## Supplementary material for "The urinary microbiome in association with diabetes and diabetic kidney disease: A systematic review": S1-S6 Tables

**S1 Table: Details of the databases and search strategy used in this systematic review.**

| **Database** | **Strategy** | **Search terms** | **Search strategy** |
| --- | --- | --- | --- |
| COCHRANE CENTRAL REGISTER OF CONTROLLED TRIALS | ‘TITLE-ABS-KEY’ | For urinary microbiome or urobiome:  ("urobiome" OR "urinary microbiome" OR "urinary microbiota" OR "urinary tract microb*" OR "urine microb*" OR "urogenital microbiome" OR "urogenital microbiota")  For diabetes, diabetic (chronic) kidney or renal disease or diabetic nephropathy:  ("kidney disease*" OR "chronic kidney disease" OR "diabetes" OR "albuminuria" OR "diabetic nephropathy" OR "end stage renal disease*" OR "renal disease*") | [  ("urobiome" OR "urinary microbiome" OR "urinary microbiota" OR "urinary tract microb*" OR "urine microb*" OR "urogenital microbiome" OR "urogenital microbiota")  AND  ("kidney disease*" OR "chronic kidney disease" OR "diabetes" OR "albuminuria" OR "diabetic nephropathy" OR "end stage renal disease*" OR "renal disease*")  ] |
| SCOPUS | ‘TITLE-ABS-KEY’ |  |  |
| MEDLINE-Ovid | ‘Keyword’ in advanced search |  |  |
| EMBASE | ‘Keyword’ in advanced search |  |  |
| PUBMED | **‘**ALL FIELDS’ |  |  |
| WEB OF SCIENCE | ‘ALL FIELDS’ |  |  |

**S2 Table: Main characteristics of selected studies**. MSU = midstream urine; MMSU = modified midstream urine; * = sequencing of microbiota extracellular vesicles; ¥ = primers for phylum Firmicutes; and genera *Bacteroides* and *Bifidobacterium.* ♀ = only females; ♂ = only males.

| **Article** | **Title** | **Study design** | **Year** | **Collection method / type of sample** [1] | **Sequen-cing method** | **Exclusion criteria** | **Country** | **Total partici-pants** | **Healthy vs diabetic adults** | **Comments** |
| --- | --- | --- | --- | --- | --- | --- | --- | --- | --- | --- |
| [2] | Impact of coexisting type 2 diabetes mellitus on the urinary microbiota of kidney stone patients | Cross-sectional case-control | 2024 | Catheteri-sed urine /  Urinary bladder | V3-V4 16S rDNA | Patients who were pregnant, menstruating, diagnosed with malignant tumors, autoimmune diseases, urethritis, prostatitis, benign prostatic hyperplasia, renal cysts, bladder inflammation, urinary abnormalities, and urinary catheterization, or used antibiotics or probiotic products within the past 4 weeks. | China | 30 | 0 vs 15 | Urinary microbiota in the renal pelvis of 15 patients with kidney stones plus T2DM and 15 patients with kidney stones alone; all of them had kidney stones as comorbidity |
| [3] | In-depth microbiological characterization of urine from subjects with type 2 diabetes | Cross-sectionalcase-control | 2024 | Voided MSU/ Urogenital | V3-V4 16S rDNA | Systemic diseases (including cancer) requiring anti-inflammatory or immunosuppressive therapies; treatment with SGLT2i or DPP4i in the previous 4 weeks, to avoid the risk of qualitative/quantitative variations of urinary bacterial populations; irritative and/or obstructive urinary or genital symptoms; current or 4 weeks prior antibiotic treatment; anatomical or functional abnormalities of the urinary tract. | Italy | 75 | 25 vs 50 | 50 patients with diabetes and no urinary symptoms, and 25 healthy controls |
| [4] | The urogenital microbiome in chronic kidney disease patients on peritoneal dialysis | Cross-sectional | 2024 | MSU / Urogenital | V3-V4 16S rDNA | Age under 18 years old; inability to consent; history of infection and /or antibiotic intake in the last three months. | Portugal | 46 | 0 vs 14 | All 46 individuals on peritoneal dialysis; 32 with other comorbidities |
| [5] | Pyuria is associated with dysbiosis of the urinary microbiota in type 2 diabetes patients receiving sodium–glucose cotransporter 2 inhibitors | Cross-sectional | 2023 | MSU / Urogenital | V3-V4 16S rDNA | Patients who were currently menstruating, using antibiotics for more than 2 weeks, or had  an indwelling catheter, all of which would likely interfere with the urinary microbiota. | Taiwan | 7 | 0 vs 7 | 7 T2DM individuals with SGLT2 treatment; 3 with pyuria in urine; 4 without |
| [6] | Urinary microbiota and serum metabolite analysis in patients with diabetic kidney disease | Cross-sectional case-control | 2023 | Voided MSU/ Urogenital | V3-V4 16S rDNA | Lack of clinical data, patients with type I diabetic nephropathy and other special types of diabetic nephropathy, combined with acute and chronic infectious diseases, acute cardiovascular and cerebrovascular diseases, antibiotics, probiotics or corticosteroid having been taken 3 months prior to sample collection. | China | 34 | 15 vs 19 | Individuals with diabetes had diabetic kidney disease. |
| [7] | Role of an unclassified *Lachnospiraceae* in the pathogenesis of type 2 diabetes: a longitudinal study of the urine microbiome and metabolites | Prospec-tive longitu-dinal and multi-center | 2022 | MSU / Urogenital | V3-V4 16S rDNA* |  | South Korea | 691 | 328 vs 199 | Also, 164 individuals with prediabetes.  Of the T2DM individuals there were 164 in the T2DM group and 35 diabetic unmatched subjects |
| [8] | Bladder microbiota are associated with clinical conditions that extend beyond the urinary tract | Cross-sectional | 2022 | Catheteri-sed urine /  Urinary bladder | V4 16S rDNA | Positive urine culture preoperatively; foreign body in bladder (e.g., indwelling catheters, ureteric stents, or bladder stones) and antibiotic treatment six weeks before. | Czech Republic | 58 ♂ | 0 vs 14 | 36 individuals did not have diabetes and 8 were unknown |
| [9] | Biochemical analysis of microbiotas obtained from healthy, prediabetic, type 2 diabetes, and obese individuals | Cross-sectional case-control | 2022 | MSU / Urogenital | 16S rDNA ¥ | Inflammatory bowel diseases and colorectal carcinoma, chronic diseases other than obesity and diabetes, pregnant and breastfeeding individuals. Medical treatment including antibiotics and oral contraceptive treatment, alcohol consumption and the absence of gastrointestinal disease and bowel related operations in the last 3 months. | Turkey | 60 | 15 vs 15 | Apart they studied 15 prediabetic and 15 obese individuals |
| [10] | Moderation effects of food intake on the relationship between urinary microbiota and urinary interleukin-8 in female type 2 diabetic patients | Cross-sectional case-control | 2020 | MMSU/ Urogenital | V3-V4 16S rDNA | UTI in the previous month; use of antibiotics, probiotics, prebiotics, or synbiotics in the previous 3 months; inability to complete the questionnaire; menstruation; urinary incontinence; anatomic urinary tract abnormalities or urinary catheter use. | China | 140 ♀ | 70 vs 70 | Same data and individuals than in [11] |
| [12] | Characteristics of the microbiota in the urine of women with type 2 diabetes | Cross-sectional case-control | 2020 | Voided MSU/ Urogenital | V4 16S rDNA | 1) Current abuse disorders, 2) bipolar depression or psychotic  disorder, 3) chronic illness, 4) severe complications of diabetes, 5) malabsorption, 6) elevated serum calcium, 7) taking St. John's Wort, 8) use of vitamin D supplements, 9) pregnant, 10) baseline systolic blood pressure 160mmHg or diastolic 100mmHg. | United States | 136 ♀ | 49 vs 87 | A genus was dominant if it had ≥ 50% of relative abundance. Urotypes defined by hierarchical clustering using Bray-Curtis dissimilarity. |
| [13] | Characteristics of urinary microflora in women with type 2 diabetic peripheral neuropathy without lower urinary tract symptoms | Cross-sectional | 2020 | MSU / Urogenital | V4 16S rDNA | 1) Antibiotic use in the past 3 months; 2) evidence of urinary tract infection in the past month; 3) There are underlying diseases that may affect urination; 4) unable to cooperate. | China | 30 ♀ | 0 vs 30 | T2DM individuals were divided: 17 with diabetic peripheral neuropathy and 13 without |
| [14] | Relationship between alterations of urinary microbiota and cultured negative lower urinary tract symptoms in female type 2 diabetes patients | Cross-sectional case-control | 2019 | MSU / Urogenital | V3-V4 16S rDNA | 1) Medical conditions that could interfere with voiding function; 2) neurological diseases; 3) urinary symptoms caused by drug abuse, spinal injury, hysterectomy, major pelvis surgery, vaginal prolapse; 4) Patients during the menstrual period; 5) recent antibiotics usage or indwelling catheter. | China | 58 ♀ | 26 vs 32 | Individuals were also assessed for lower urinary tract symptoms. |
| [15] | Diversity of the midstream urine microbiome in adults with chronic kidney disease | Cross-sectional | 2018 | Voided MSU/ Urogenital | V4 16S rDNA | 1) Immunosuppression medication, 2) history of a neobladder or ileal conduit, 3) had used antibiotics  within the past 4 weeks, 4) had active cancer treatment, or 5) had urinary instrumentation within the past 6 months. | United States | 77 | 23 vs 54 | Dominant urotype defined as > 50% of sequences. |
| [16] | Characterization of the urinary microbiota of elderly women and the effects of type 2 diabetes and urinary tract infections on the microbiota | Cross-sectional case-control | 2017 | MMSU/  Urogenital | V3-V4 16S rDNA | UTIs in the previous month; use of antibiotics, probiotics, prebiotics, or synbiotics in the previous 3 months; unable to complete the questionnaire; menstruation; urinary incontinence; known anatomic urinary tract abnormalities; urinary catheter. | China | 100 ♀ | 50 vs 50 | Elderly vs non-elderly cohorts. Half of the females of each cohort had T2DM |
| [11] | Dysbiosis of urinary microbiota is positively correlated with Type 2 diabetes mellitus | Cross-sectional case-control | 2017 | MMSU/ Urogenital | V3-V4 16S rDNA | UTIs in the previous month; use of antibiotics, probiotics, prebiotics, or synbiotics in the previous 3 months; unable to complete the questionnaire; menstruation; urinary incontinence; known anatomic urinary tract abnormalities; urinary catheter. | China | 140 ♀ | 70 vs 70 | The same cohort data was also analysed in [10]; in [17]; and [18] |
| [17] | Alterations of urinary microbiota in type 2 diabetes mellitus with hypertension and/or hyperlipidaemia | Cross-sectional case-control | 2017 | MMSU/ Urogenital | V3-V4 16S rDNA | UTIs in the previous month; use of antibiotics, probiotics, prebiotics, or synbiotics in the previous 3 months; unable to complete the questionnaire; menstruation; urinary incontinence; known anatomic urinary tract abnormalities; urinary catheter. | China | 70 ♀ | 0 vs 70 | Same diabetic cohort than in previous article [11]. Cohort divided in 28 T2DM individuals; 24 T2DM and hypertension; 7 T2DM and hyperlipidaemia; 11 T2DM and hypertension and hyperlipidaemia. |
| [18] | Dysbiosis of the urinary microbiota associated with urine levels of proinflammatory chemokine interleukin-8 in female type 2 diabetic patients | Cross-sectional case-control | 2017 | MMSU/ Urogenital | V3-V4 16S rDNA | UTIs in the previous month; use of antibiotics, probiotics, prebiotics, or synbiotics in the previous 3 months; unable to complete the questionnaire; menstruation; urinary incontinence; known anatomic urinary tract abnormalities; urinary catheter. | China | 70 ♀ | 0 vs 70 | Same diabetic cohort than in [11] |
| [19] | Urinary microbiome of kidney transplant patients reveals dysbiosis with potential for antibiotic resistance | Cross-sectional case-control | 2017 | MSU / Urogenital | Shotgun | Control group without pre-existing medical conditions such as kidney disease, UTI or others; not consume any antibiotics (for at least 6 months) prior to participation in the study and not report any history of over-the-counter medications such as antipyretics or pain-relievers. | United States | 29 | 8 vs 6 | Individuals with diabetic renal disease (type 1 and type 2 diabetes). But also, 15 individuals had non-diabetic renal disease as there were different primary diagnoses for renal disease. |

**S3 Table: Quality and risk of bias assessment of selected studies based on Cochrane and NHLBI** [20,21]. Y: yes; N: no; CD: cannot determine; NR: not reported. ^£^ = sequencing of microbiota extracellular vesicles; ^¥^ = primers for phylum Firmicutes; and genera *Bacteroides* and *Bifidobacterium;* ^Ω^ = some or all individuals used antibiotic prophylaxis. ♀ = only females; ♂ = only males.

| Article | Q1 | Q2 | Q3 | Q4 | Q5 | Q6 | Q7 | Q8 | Q9 | Q10 | Q11 | Risk of bias | (If applicable) Explanation for high risk |
| --- | --- | --- | --- | --- | --- | --- | --- | --- | --- | --- | --- | --- | --- |
| [2] | Y | Y | Y | Y | Y | Y | CD | Y | Y | NR | Y | Low |  |
| [3] | Y | Y | NR | Y | Y | Y | CD | Y | Y | NR | N | High | No statistical comparison between clinical features of controls and diabetic individuals that may affect the microbiota (e.g. BMI) only between males and females in each of the groups. |
| [4] | Y | Y | Y | Y | Y | Y | CD | Y | Y | NR | Y | Low |  |
| [5] | Y | Y | Y | Y | Y | Y | CD | Y | Y | NR | Y | Low |  |
| [6] | Y | Y | NR | Y | Y | Y | CD | Y | Y | NR | Y | Low |  |
| [7] ^£^ | Y | Y | NR | Y | NR | Y | CD | Y | Y | NR | Y | Low |  |
| [8] ♂ ^Ω^ | Y | Y | Y | Y | Y | Y | CD | Y | Y | NR | Y | Low |  |
| [9] ^¥^ | Y | Y | NR | Y | Y | Y | CD | Y | Y | NR | Y | Low |  |
| [10] *♀ | Y | Y | NR | Y | Y | Y | CD | Y | Y | NR | Y | Low |  |
| [12] ♀ | Y | Y | Y | Y | Y | Y | CD | Y | Y | NR | Y | Low |  |
| [13] ♀ | Y | Y | Y | Y | Y | Y | CD | Y | Y | NR | Y | Low |  |
| [14] ♀ | Y | Y | Y | Y | Y | Y | CD | Y | Y | NR | Y | Low |  |
| [15] | Y | Y | Y | Y | Y | Y | CD | Y | Y | NR | Y | Low |  |
| [16] ♀ | Y | Y | NR | Y | Y | Y | CD | Y | Y | NR | Y | Low |  |
| [11] ♀ | Y | Y | NR | Y | Y | Y | CD | Y | Y | NR | Y | Low |  |
| [17]* ♀ | Y | Y | NR | Y | Y | Y | CD | Y | Y | NR | Y | Low |  |
| [18] *♀ | Y | Y | NR | Y | Y | Y | CD | Y | Y | NR | Y | Low |  |
| [19] ^Ω^ | Y | Y | Y | Y | Y | Y | CD | Y | Y | NR | N | High | Patients, but not controls, received prophylactic antibiotic which affects diversity. They considered folate pathways to check the impact of that (target of TMP/SXT). |

Q1. Was the research question or objective in this paper clearly stated and appropriate?
Q2. Was the study population clearly specified and defined?
Q3. Did the authors include a sample size justification?
Q4. Were controls selected or recruited from the same or similar population that gave rise to the cases (including the same timeframe)?
Q5. Were the definitions, inclusion/exclusion criteria, algorithms or processes used to identify or select cases and controls valid, reliable, and implemented consistently across all study participants?
Q6. Were the cases clearly defined and differentiated from controls?
Q7. If less than 100 percent of eligible cases and/or controls were selected for the study, were the cases and/or controls randomly selected from those eligible?
Q8. Were the investigators able to confirm that the disease occurred prior to the development of the condition or event that defined a participant as a case?
Q9. Were the measures of diseases vs without clearly defined, valid, reliable, and implemented consistently (including the same period) across all study participants?
Q10. Were the assessors of diseases vs without blinded to the case or control status of participants?
Q11. Were key potential confounding variables measured and adjusted statistically in the analyses? If matching was used, did the investigators account for matching during study analysis?

**S4 Table: Specific diversity, evenness and richness indexes used in selected studies**.

| **Reference** | **Title** | **Number of OTUS** | **Observed species** | **Chao** | **ACE** | **Shannon** | **Simpson** | **Inverse Simpson** |
| --- | --- | --- | --- | --- | --- | --- | --- | --- |
| [2] | Impact of coexisting type 2 diabetes mellitus on the urinary microbiota of kidney stone patients |  |  | X |  | X |  |  |
| [3] | In-depth microbiological characterization of urine from subjects with type 2 diabetes |  | X | X |  | X |  |  |
| [4] | The urogenital microbiome in chronic kidney disease patients on peritoneal dialysis |  |  |  |  | X |  |  |
| [5] | Pyuria is associated with dysbiosis of the urinary microbiota in Type 2 diabetes patients receiving sodium–glucose cotransporter 2 inhibitors |  |  |  |  | X |  |  |
| [6] | Urinary microbiota and serum metabolite analysis in patients with diabetic kidney disease |  |  | X | X | X | X |  |
| [7] | Role of an unclassified Lachnospiraceae in the pathogenesis of type 2 diabetes: a longitudinal study of the urine microbiome and metabolites |  |  | X | X | X | X |  |
| [8] | Bladder microbiota are associated with clinical conditions that extend beyond the urinary tract |  |  | X | X | X | X |  |
| [9] | Biochemical analysis of microbiotas obtained from healthy, prediabetic, type 2 diabetes, and obese individuals |  |  |  |  |  |  |  |
| [10] | Moderation effects of food intake on the relationship between urinary microbiota and urinary interleukin-8 in female type 2 diabetic patients | X | X | X | X | X | X |  |
| [12] | Characteristics of the microbiota in the urine of women with type 2 diabetes |  |  |  |  |  |  |  |
| [13] | Characteristics of urinary microflora in women with type 2 diabetic peripheral neuropathy without lower urinary tract symptoms; [无下尿路症状的女性2型糖尿病周围神经病变患者的尿液菌群特征] | X | X | X | X | X | X |  |
| [14] | Relationship between alterations of urinary microbiota and cultured negative lower urinary tract symptoms in female type 2 diabetes patients | X | X | X | X | X | X |  |
| [15] | Diversity of the midstream urine microbiome in adults with chronic kidney disease |  |  | X |  | X |  | X |
| [16] | Characterization of the urinary microbiota of elderly women and the effects of type 2 diabetes and urinary tract infections on the microbiota | X |  | X |  | X | X |  |
| [11] | Dysbiosis of urinary microbiota is positively correlated with Type 2 diabetes mellitus | X | X | X | X | X | X |  |
| [17] | Alterations of urinary microbiota in type 2 diabetes mellitus with hypertension and/or hyperlipidaemia | X | X | X | X | X | X |  |
| [18] | Dysbiosis of the urinary microbiota associated with urine levels of proinflammatory chemokine interleukin-8 in female type 2 diabetic patients | X | X | X | X | X | X |  |
| [19] | Urinary microbiome of kidney transplant patients reveals dysbiosis with potential for antibiotic resistance. |  | X |  |  | X |  | X |

**S5 Table: Characteristics and results of selected studies.** BMI = body mass index; CKD = chronic kidney disease; DKD = diabetic kidney disease; DPN = diabetic peripheral neuropathy; eGFR = estimated glomerular filtration rate; ESRD = end stage renal disease; FBG = fasting blood glucose; H = healthy controls; HbA1c = glycosylated haemoglobin; IL-8= interleukin 8; NS = Not stated; OTU = operational taxonomic unit; PD = peritoneal dialysis; SBP = systolic blood pressure; (TX)DM = (type X = 1 or X = 2) Diabetes mellitus; SD = standard deviation; vs = versus; * = studies that used data from participants/cohort of article 13 authored by Liu et al. 2017 [11]; ^£^ = sequencing of microbiota extracellular vesicles; ^¥^ = primers for phylum Firmicutes; and genera *Bacteroides* and *Bifidobacterium;* ^Ω^ = some or all individuals used antibiotic prophylaxis. ♀ = only females; ♂ = only males; *↓* = lower; ↑ = higher.

| **Article** | | **Mean age ± SD** | **Context** | **Indexes**  **results in diabetic individuals** | **Main findings in relative abundance of diabetic individuals in urine [if stated, differences in prevalence]** | **Other significant details or relevant information** |
| --- | --- | --- | --- | --- | --- | --- |
| Case-control studies with controls and diabetic individuals | | | | | | |
| [2] | | Kidney stones and T2DM:  56 ± 11  Kidney stones alone:  55 ± 11 | All individuals with kidney stones. Half of them had also T2DM. | No differences in richness (Chao index)  ↑ α - diversity (Shannon index)  β-diversity: the group with kidney stones alone and with kidney stones and T2DM clustered and could be differentiated | ↑*Sphingomonas*  ↑*Propionibacterium*  ↑ *Corynebacterium*  ↑ *Cellulosimicrobium*  ↑ *Methylophilus*  ↑ *Lactobacillus*  ↑ *Enhydrobacter*  ↑ *Chryseobacterium*  ↑ *Haemophilus*  ↑ *Allobaculum* | The relative abundance of *Enhydrobacter, Chryseobacterium,* and *Allobaculum*  genera exhibited correlations with fasting blood glucose and HbA1c values. |
| [3] | | H:  62 ± 7 (males)  57 ± 6 (females)  T2DM:  65 ± 9 (males)  64 ± 10 (females) |  | No differences in richness or α - diversity (observed species, Chao and Shannon indexes) | NS  [Statistical differences in prevalence (males and females):  *Bifidobacterium breve, Corynebacterium pyroviciproducens, Lactobacillus gasseri ,* and *Streptococcus agalactiae* were more prevalent in healthy controls. *Campylobacter ureolyticus, Corynebacterium coyleae, Corynebacterium glucuronolyticum, Enterococcus faecalis, Escherichia coli, Lactobacillus iners, Peptoniphilus grossensis,* and *Veillonella atypica* were more prevalent in T2DM individuals.  *Actinotignum schaalii, Bifidobacterium scardovii, Facklamia hominis, Negativicoccus succinicivorans,* and *Peptoniphilus lacrimalis* were less prevalent in T2DM males compared to healthy male controls. *Aerococcus christensenii, Anaerococcus hydrogenalis, Brevibacterium ravenspurgense, Corynebacterium aurimucosum, Gardnerella vaginalis, Mobiluncus curtisii, Prevotella buccalis, Prevotella colorans*, and *Veillonella montpellierensis* were more prevalent in T2DM females whereas *Facklamia ignava* and *Winkia neuii* were less prevalent in T2DM females compared to female healthy controls.] | Total bacterial load and the abundance of total Bacillota were found to be elevated in patients with diabetes compared with the healthy control group, this was also true for females but not males.  *Actinotignum urinale, Aerococcus christensenii , Corynebacterium coyleae, Enterococcus faecalis , Escherichia coli , Ezakiella massiliensis, Gardnerella vaginalis, Peptoniphilus grossensis, Prevotella amnii* and *Veillonella montpellierensis* were only detectable in T2DM individuals *while Bifidobacterium scardovii* was the only species not found in patients with T2DM.  *Actinobaculum massiliense, Actinotignum urinale, Aerococcus christensenii , Anaerococcus hydrogenalis, Campylobacter ureolyticus, Corynebacterium aurimucosum, Corynebacterium coyleae, Corynebacterium glucuronolyticum, Enterococcus faecalis , Escherichia coli , Ezakiella massiliensis, Gardnerella vaginalis, Lactobacillus iners, Mobiluncus curtisii, Peptoniphilus grossensis, Porphyromonas uenonis, Prevotella amnii, Prevotella buccalis, Veillonella atypica,* and *Veillonella montpellierensis* were significantly more prevalent in patients with T2DM;  *Actinotignum schaalii, Bifidobacterium breve*, *Bifidobacterium longum, Bifidobacterium scardovii, Corynebacterium pyruviciproducens, Dialister propionifaciens, Facklamia hominis, Lactobacillus gasseri, Negativicoccus succinicivorans, Peptoniphilus lacrimalis, Schaalia radingae, Streptococcus agalactiae, Streptococcus anginosus* , and *Winkia neuii* were significantly more prevalent in healthy controls. |
| [6] | | H: 44 ± 8 T2DM:  60 ± 9 | T2DM with DKD | *↓* α - diversity  in T2DM  (Shannon and Simpson indexes) | *↓*Bacillota (Firmicutes)  *↓* Bacteroidota *(Bacteroidetes)*  ↑ Pseudomonadota (Proteobacteria)  ↑Acidobacteriota (Acidobacteria) | H and T2DM clinical characteristics differed significantly in age; BMI; SBP; urea; uric acid; creatinine; natriuretic peptide; HbA1c (all ↓in H); and HDL and albumin (both ↑ in H).  HbA1c and metabolites related to arginine and proline were positively correlated with Acidobacteria.  eGFR was positively correlated with Bacillota, Clostridiales, Clostridia, *Lactobacillaceae* and *Lactobacillus*. |
| [7] ^£^ | | 59 ± 6  (average of all groups and periods) | Longitudinal changes over 4 years sequencing extracellular vesicles of H, prediabetics and T2DM individuals. | *↓* α - diversity  (Shannon index) in all groups  (possible effect of aging) | *↓* unclassified *Lachnospiraceae* GU174097_g | Low abundance of GU174097_g was a risk factor for T2DM development, and it was associated with T2DM progression. GU174097_g decreased ketone bodies levels; and thus reduced HbA1c levels. |
| [8] ♂ ^Ω^ | | 65 ± 14 | Catheterized urine samples from men collected under anesthesia from the bladder prior to urological surgery | *↓* richness  (OTUs, ACE and iChao2 indexes)  *↓* α - diversity  (Shannon and Simpson indexes)  β - diversity  was no different | No statistically differences of specific taxa between T2DM and non-T2DM individuals | Lower richness was also associated with: (1) ≥75 years (index iChao2); (2) High cholesterol and/or hyperlipidemia (ACE and iChao2 indexes); (3) antibiotic prophylaxis (OTUs and iChao2)  Current smokers had higher α - diversity (Simpson index).  No statistically significant differences in α - diversity for hypertension or CKD.  β – diversity (Bray-Curtis dissimilarity) was different between individuals with (1) CKD vs no CKD; (2) severe vs mild urinary symptoms; and (3) antibiotic prophylaxis vs without prophylaxis |
| [9] ^¥^ | | H: 42 ± 11  Obese:  42 ± 9  Pre-DM:  50 ± 9  T2DM: 54 ± 8 | T2DM, prediabetic, obese and HC | NS | *↓ Bifidobacterium* (same was observed for obese and prediabetics) | Clinical features that were statistically different: FBG between all groups; HbA1c between all groups except for H and obese individuals; BMI between all groups except for prediabetics and both obese and T2DM. Age was significantly lower in both H and obese vs T2DM individuals. Triglycerides and HDL were significantly lower in H vs both prediabetics and T2DM individuals |
| [12] ♀ | | H: 51 ± 11  T2DM: 51 ± 11 | Females with T2DM | β – diversity:  Both H and T2DM clustered in 4-5 urotypes    Urotypes in T2DM:  *Lactobacillus* and *Enterobacteriaceae* vs  H females: *Gardnerella* and mixed | ↑ *Lactobacillus*  ↓ *Corynebacterium*  ↓ *Staphylococcus*  [Statistical differences in presence/absence:  *Lactobacillus* was more present while *Corynebacterium, Anaerococcus, Finegoldia* and *Peptoniphilus* were less present in T2DM] | H and T2DM group clinical characteristics differed significantly in FBG and HbA1c (both ↓in H). HbA1c was positively correlated with *Lactobacillus* and negatively associated with *Prevotella* and *Corynebacterium* in all cohort; and negatively associated with *Prevotella* also in the T2DM group.  Prevalence of *Lactobacillus* increased with weight; but BMI and urotypes associations were non-significant |
| [14] ♀ | | H: 58 ± 9  T2DM:  57 ± 8 | Females H and T2DM. T2DM microbiota were also assessed for urinary symptoms and levels of HbA1c | Richness and α – diversity were no different (OTUs, observed species, Chao, ACE, Shannon and Simpson indexes)  β - diversity  was different between H and T2DM | ↑ *Escherichia-Shigella*  ↑ *Klebsiella*  ↑ *Aerococcus*  ↑*Delftia*  ↑*Enterococcus*  ↑*Alistipes*  ↑*Stenotrophomonas*  ↑ *Micrococcus*  ↑ *Deinococcus*  ↑ *Rubellimicrobium*  ↓ *Gallicola*  ↓*Arcobacter*  ↓*Arcanobacterium*  ↓*Kocuria*  ↓ *Murdochiella*  ↓ *Solitalea*  ↓ *Peptoniphilus* | The clinical feature that was statistically different between H and T2DM individuals was FBG (FBG ↑ in T2DM).  The clinical feature that was statistically different between T2DM individuals with urinary symptoms and without was HbA1c (HbA1c ↑ in T2DM with urinary symptoms).  The diabetic group with urinary symptoms had lower richness and different β – diversity but no differences in α – diversity compared to the diabetic group without urinary symptoms.    ↑*Escherichia-Shigella*  ↑ *Campylobacter*  ↑ *Megasphaera*  ↓ *Prevotella*  ↓ *Dialister*  ↓ *Leptotrichia*  ↓ *Anaerococcus*  ↓ *Fusobacterium*  ↓ *Prevotella_6,*  ↓ *Fastidiosipila*  ↓ *Varibaculum,*  ↓ *Mycoplasma*  ↓ *Peptoniphilus*  ↓ *Porphyromonas*  ↓ *Fodinicola*  ↓ *Brevundimonas*  ↓ *Candidatus- Nomurabacteria*  ↓ *Paracoccus*  ↓ *Rhodococcus*  The diabetic group with high HbA1c had lower α – diversity and different β – diversity but no difference in richness compared to the diabetic group with low HbA1c.  ↑*Escherichia-Shigella*  ↑ *Lactobacillus*  ↓ *Prevotella*  ↓ *Campylobacter*  ↓ *Dialister*  ↓ *Anaerococcus*  ↓ *Peptoniphilus* ↓ *Porphyromonas*  ↓ *Fodinicola*  ↓ *Negativicoccus* |
| [16] ♀ | | Elderly:  72 ± 7  No-old:  50 ± 8 | Old and non-old cohorts where half of the females in each one had T2DM. Comparison between age and T2DM status. | *↓* richness  (OTU and Chao indexes) in elderly with T2DM | Elderly with T2DM (compared to elderly without):  *↓ Nitrospirae*  *↓* *Odoribacter*  *↓Aeromonas*  *↓Clostridium*  *↓Agrobacterium*  *↓Desulfovibrio*  *↓Bilophila*  *↓Enterobacter*  *↓Butyricimonas*  *↓Enterococcus*  *↓ Erwinia*  *↓ Fusobacterium*  *↓ Klebsiella*  *↓ Lachnobacterium*  *↓Lysobacter*  *↓Mitsuokella*  *↓Stenotrophomonas*  *↓Rhodoplanes*  *↓Ramlibacter*  *↓Phascolarctobacterium*  ↑*Eggerthella*  ↑ *Parvimonas*  ↑*Turicibacter*  ↑*Bdellovibrio*  ↑*Modestobacter*  ↑*Hydrogenphaga*  ↑*Anaeromyxobacter*  ↑ *Microbacterium*  ↑***Lactobacillus iners*** | T2DM females treated with metformin. *Lactobacillus* had a decreased trend in T2DM patients with FBG > 10 mmol/l. But *Lactobacillus iners* was not associated with FBG.  Elderly related differences: (1) relative abundance of Bacillota (Firmicutes) increased with BMI; (2) relative abundance of Bifidobacteria was negatively correlated with age; (3) *Lactobacillus* decreased with age but was not associated with pH; (4) higher abundance of *Sphingobium* and *Bosea;* and (5) lower abundance of *Sneathia; Geobacillus; Shuttleworthia; Bacillus; Gemella; Bdellovibrio; Hydrogenphaga; Proteus; Novosphingobium and Cateribacterium*. |
| [11] ♀ | | Matched H and T2DM individuals.  70% > 56 years or older; only 8% younger than 45 years | Authors matched individual characteristics between the H and the T2DM groups | ↓ richness  (number of OTUs, observed species, ACE and Chao1 indexes)    ↓ α-diversity (Shannon index) | ↑ Actinobacteria  ↑ *Porphyromonas*  ↑ *Flavobacteriales*  ↑ *Flavobacteria*  ↑ *Collinsella*  *↓ Akkermansia muciniphila*  *↓* Bacteroidia  *↓* Bacteroidetes  *↓* *Prevotellaceae*  *↓ Prevotella*  *↓Pseudomonadales*  *↓Pseudomonas*  *↓Peptoniphilus*  *↓ Citrobacter*  *↓*Actinobacteria  *↓ Synergistales*  *↓ Synergistetes*  *↓Acinetobacter*  *↓Campylobacter*  *↓Campylobacterales*  *↓ Staphylococcus*  *↓ Anaerococcus*  *↓ Halomonas*  *↓ Moraxellaceae*  *↓ Finegoldia*  *↓ Streptococcus*  *↓Veillonella*  *↓Clostridium*  *↓Corynebacterium* | Actinobacteria increase was associated with higher BMI; increased FBG and urine glucose.  *Akkermansia muciniphila* decrease was associated with FBG and urine glucose.  Carbohydrate and amino acid metabolism was damaged in T2DM patients, and correlated with bacterial diversity.  Clinical features statistically different between H and T2DM individuals were, urine glucose, FBG, urine infections, hypertension and hyperlipidemia ( all ↑ in T2DM). Authors studied specifically hypertension and hyperlipidemia in article 14 below [17]; inflammatory markers in article 15 (Ling et al. 2017) and the effect on diet in article 7 [10]. |
| [19] ^Ω^ | | H: 35  ESDR T1DM: 51  ESDR T2DM: 64  (all ESRD: 53) | Kidney transplant ESRD patients caused by four primary diagnoses: T1DM, T2DM, hypertension and ‘other’.  Patients received trimethoprim -sulfamethoxa-zole. | ↓ α-diversity (Shannon and inverse Simpson indexes)  ↓ richness (observed species)  β-diversity: primary diagnoses had no significant differences in bacterial composition; but all had with the H | ↑ *Enterococcus faecalis*  ↑ *Enterococcus faecium*  ↑ *Enterococcus sp*.  (↑ Bacillota)  ↑ *Escherichia coli*  ↑ *Escherichia sp.*  ↓ *Cutibacterium acnes*  ↓ *Corynebacterium*  ↓*Mobiluncus curtisii*  (↓Actinobacteria) | Prophylactic antibiotic: both groups had similar folate metabolism (which is the target of trimethoprim -sulfamethoxazole) but differences in specific enzymes. Dihydrofolate synthase/folypolyglutamate synthase was increased in transplant groups. |
| Studies with diabetic individuals only | | | | | | |
| [5] | | T2DM No-pyuria: 59 ± 6  T2DM Pyuria: 60 ± 7 | All individuals with T2DM (SGLT2  therapy) | ↓ α - diversity  in T2DM with pyuria compared with T2DM without pyuria (Shannon index) | ↑*Escherichia-Shigella* in individuals with pyuria | 7 T2DM individuals with SGLT2 treatment; all with negative standard urine cultures. eGFR stage II (60–89 mL/min) mildly impaired kidney function.  *Staphylococcus* was positively correlated with HDL. |
| [13] ♀ | | T2DM DPN:  56 ± 8  T2DM only:  55 ± 7 | Females with T2DM with DPN and without DPN | ↓richness in DPN group (OTUs, observed species, ACE and iChao2 indexes)  α – diversity was no different  (Shannon and Simpson indexes)  β - diversity  was no different | Group with DPN had:  ↑ *Mycoplasmataceae ↓*Propionibacteriaceae  ↓ Pseudobdellovibrioinaceae ↓Sphingobacteriaceae | T2DM females: 17 with DPN and 13 without.  Both groups did not differ in clinical characteristics. |
| *♀ | [17] | T2DM:  56 ± 14  T2DM + hypertension:  70 ± 9  T2DM + hyperlipidemia:  54± 11  T2DM + both: 70 ± 10 | Only analysis of the T2DM individuals from article 13 that had different clinical features: hypertension; hyperlipidemia, none or both. | ↑ richness compared to the hyperlipidemia only group (number of reads and number of OTUs)  ↓ richness compared to hypertension groups (number of reads and number of OTUs) | Compared to T2DM + hypertension group, the group **with only** T2DM had:  ↑ *Lactobacillus iners*  ↑ *Acinetobacter rhizosphaerae*  ↑ *Acinetobacter schindleri*  ↑*Lactobacillus*  ↑*Enterobacter*  ↑*Klebsiella*  ↑*Shuttleworthia*  ↑*Sneathia*  ↑ *Parvimonas*  ↑*Megasphera*  ↑*Erwinia*  *↓ Kocuria palustris*  *↓ Aeromonas*  *↓ Roseburia*  *↓ Ruminococcus*  Compared to T2DM + hyperlipidemia, the group **with only** T2DM had:  *↓ Faecalibacterium*  *↓ Collinsella*  *↓ Oscillospira* ( that genus was a diagnostic factor to differentiate both groups)  Compared to T2DM + hypertension and hyperlipidemia, the group **with only** T2DM had:  ↑*Acinetobacter rhizosphaerae*  ↑ *Shuttleworthia*  *↓Gemella*  *↓Prevotella*  Compared to T2DM + hypertension and hyperlipidemia, the group with T2DM + hyperlipidemia had:  ↑*Acinetobacter rhizosphaerae*  ↑*Faecalibacterium*  Compared to T2DM + hypertension, the group with T2DM + hyperlipidemia had:  ↑ *Klebsiella*  ↑ *Novosphingobium*  ↑ *Lactobacillus*  ↑ *Enterobacter*  The two groups with hypertension did not show significant differences | The T2DM cohort was divided: 28 T2DM; 24 T2DM + hypertension; 7 T2DM + hyperlipidemia; 11 T2DM + hypertension + hyperlipidemia. The duration of T2DM was significantly shorter in the two groups without hypertension.  The four most abundant bacteria in all groups were Proteobacteria, Bacillota, Bacteroidota and Actinobacteriota. The fifth varied from Fusobacteria (in T2DM and T2DM with both conditions), Synergistota (group with hypertension) and Acidobacteriota (group with hyperlipidemia).  Predominant genera: in T2DM *Lactobacillus, Prevotella,* and *Acinetobacter;* in T2DM + hyperlipidemia *Lactobacillus, Prevotella* and *Halomonas*; in T2DM + hypertension *Prevotella, Streptococcus*, and *Bacteroides*; and in T2DM + hypertension + hyperlipidemia *Prevotella, Lactobacillus* and *Bacillus*.  The groups with hypertension had presence of *Eggerthella lenta* but this species was absent in the group with only diabetes*;* while *Gardnerella* was absent in the hyperlipidemia group but it was present in the other groups.  Several bacteria correlated with FBG, blood pressure and lipidic profiles. *Atopobium* was positively correlated with FBG in the T2DM group.  They explored the association between nutrient intake and the relative abundance of bacteria in urine which is detailed in article 7 [10]. |
| *♀ | [18] | T2DM with IL-8 in urine:  66 ± 14  T2DM without IL-8 in urine:  59 ± 11 | T2DM females whose levels of IL-8 were evaluated. IL-8 is a potential biomarker for diagnosing of urinary infections | α – diversity was no different between T2DM females with and without IL-8 in urine (Shannon and Simpson indexes)  β-diversity: the group with IL-8 and without IL-8 in urine clustered and could be differentiated | T2DM females without urinary IL-8:  ↓*Bifidobacteriaceae* ↓*Shuttleworthia*  ↓ *Thermus*  ↓*Thermales*  ↓*Thermaceae*  ↓ *Streptococcus anginosus*  ↓ *Acinetobacter rhizosphaerae*  ↓ *Acinetobacter schindleri*  ↓ *Lactobacillus iners*  ↓ *Akkermansia muciniphila*  ↓ *Mobiluncus*  ↓ *Peptoniphilus*  ↓ *Corynebacterium*  ↓ *Gemella*  ↓ *Enterococcus*  ↓ *Aquaspirillum*  ↓ *Geobacillus*  ↑ *Prevotella copri*  ↑ *Prevotella stercorea*  ↑*Bacteroides uniformis* ↑*Coprococcus eutactus*  ↑ *Faecalibacterium*  ↑ *Megamonas*  ↑ *Comamonas*  ↑ *Sutterella*  ↑*Pseudomonas*  ↑*Phascolarctobacterium*  ↑*Parabacteroides*  ↑ Pseudomonadota  ↑ Bacteroidota | *Ruminococcus* was positively associated with urinary IL-8. Urinary IL-8 was low in T2DM patients with ↑ *Acinetobacter*, ↑*Microbacterium* and ↑*Megamonas*.  Urinary IL-8 was high in T2DM patients with ↑ *Pseudomonas* and ↑ *Klebsiella*.  Age, BMI, pH, white blood cells, leukocyte esterase, protein, glucose, and nitrites in urine between the T2DM groups with IL-8 and without IL-8 in differed and all were ↑ in the group with IL-8 |
| *♀ | [10] | Matched H and T2DM individuals.  70% > 56 years or older; only 8% younger than 45 years | T2DM association with urinary IL-8 and evaluation of the effect of diet on urinary microbiota | NS | NS | *Ruminococcus* was positively associated with urinary IL-8 (results from article 15 by Ling et al. 2017). Water weakened the positive relationship between *Ruminococcus* and IL-8 in urine while fiber, vitamin B3 and vitamin E enhanced the positive relationship.  Cholesterol and magnesium had a positive association between the relative abundance of *Comamonas* and IL-8 concentration in urine. |
| Case-control studies with individuals that have kidney or renal disease | | | | | | |
| [4] | | 57 ± 11 | All individuals with CKD | ↓ α - diversity (Shannon index) as CKD-PD worsened (independently of the cause) | *↓Atopobium*  *↓ Dermabacter*  *↓Gardnerella* | Apart from diabetes status in CKD patients, significance was found for sex. *Lactobacillus* linked to females and *Staphylococcus* and *Anaerococcus* to males. |
| [15] | | Males:  77 ± 8  Females:  71 ± 8 | All individuals with CKD but no dialysis (70% of them with diabetes) | ↓ α - diversity (Shannon index) as CKD worsened (independently of the cause)  β – diversity:  Urotypes in CKD (number of samples): *Corynebacterium*(11); *Staphylococcus* (9); *Lactobacillus* (7); *Gardnerella* (7); *Streptococcus* (7); P*revotella* (4); *Escherichia-Shigella* (3); Enterobacteriaceae (2) | NS | Richness and α – diversity were no different between diabetic and non-diabetic CKD individuals nor different by age or obesity status (Chao; Inverse Simpson and Shannon indexes).  Higher eGFR was associated with ↑ α-diversity  Urgent urinary incontinence was associated with ↑ α-diversity |

**S6 Table: Taxonomic classification of bacteria identified in selected studies according to GTDB release 220** (<https://gtdb.ecogenomic.org/>). N = number of articles; * Taxonomic names have been modified.

| **Bacteria** | **N** | **PHYLUM** | **CLASS** | **ORDER** | **FAMILY** | **GENUS** | **SPECIES** |
| --- | --- | --- | --- | --- | --- | --- | --- |
| *Lower abundance or less prevalent in urine of diabetics* |  |  |  |  |  |  |  |
| **Actinobacteriota** | **1** | Actinobacteriota |  |  |  |  |  |
| ***Actinotignum schaalii*** | **1** | Actinomycetota | Actinomycetes | Actinomycetales | Actinomycetaceae | Actinotignum | schaalii |
| ***Arcanobacterium*** | **1** | Actinomycetota | Actinomycetes | Actinomycetales | Actinomycetaceae | Arcanobacterium |  |
| ***Mobiluncus curtisii*** | **1** | Actinomycetota | Actinomycetes | Actinomycetales | Actinomycetaceae | Mobiluncus | curtisii |
| ***Winkia neuii*** | **1** | Actinomycetota | Actinomycetes | Actinomycetales | Actinomycetaceae | Winkia | neuii |
| ***Bifidobacterium*** | **1** | Actinomycetota | Actinomycetes | Actinomycetales | Bifidobacteriaceae | Bifidobacterium |  |
| ***Bifidobaterium breve*** | **1** | Actinomycetota | Actinomycetes | Actinomycetales | Bifidobacteriaceae | Bifidobacterium | breve |
| ***Bifidobaterium scardovii*** | **1** | Actinomycetota | Actinomycetes | Actinomycetales | Bifidobacteriaceae | Bifidobacterium | scardovii |
| ***Gardnerella**** | **1** | Actinomycetota | Actinomycetes | Actinomycetales | Bifidobacteriaceae | Bifidobacterium |  |
| ***Dermabacter*** | **1** | Actinomycetota | Actinomycetes | Actinomycetales | Dermabacteraceae | Dermabacter |  |
| ***Kocuria*** | **1** | Actinomycetota | Actinomycetes | Actinomycetales | Micrococcaceae | Kocuria |  |
| ***Corynebacterium*** | **3** | Actinomycetota | Actinomycetes | Mycobacteriales | Mycobacteriaceae | Corynebacterium |  |
| ***Corynebacterium pyroviciproducens*** | **1** | Actinomycetota | Actinomycetes | Mycobacteriales | Mycobacteriaceae | Corynebacterium | pyroviciproducens |
| ***Cutibacterium acnes*** | **1** | Actinomycetota | Actinomycetes | Propionibacteriales | Propionibacteriaceae | Cutibacterium | acnes |
| ***Atopobium*** | **1** | Actinomycetota | Coriobacteriia | Coriobacteriales | Atopobiaceae | Atopobium |  |
| **Bacillota** | **1** | Bacillota |  |  |  |  |  |
| **Flackamia hominis** | **1** | Bacillota | Bacilli | Lactobacillales | Aerococcaceae | Flackamia | hominis |
| **Flackamia ignava *** | **1** | Bacillota | Bacilli | Lactobacillales | Aerococcaceae | Falseniella | ignava |
| ***Enterococcus*** | **1** | Bacillota | Bacilli | Lactobacillales | Enterococcaceae | Enterococcus |  |
| ***Lactobacillus gasseri*** | **1** | Bacillota | Bacilli | Lactobacillales | Lactobacillaceae | Lactobacillus | gasseri |
| ***Streptococcus*** | **1** | Bacillota | Bacilli | Lactobacillales | Streptococcaceae | Streptococcus |  |
| ***Streptococcus agalactiae*** | **1** | Bacillota | Bacilli | Lactobacillales | Streptococcaceae | Streptococcus | agalactiae |
| ***Staphylococcus*** | **2** | Bacillota | Bacilli | Staphylococcales | Staphylococcaceae | Staphylococcus |  |
| ***Clostridium*** | **2** | Bacillota | Clostridia | Clostridiales | Clostridiaceae | Clostridium |  |
| ***Lachnobacterium*** | **1** | Bacillota | Clostridia | Lachnospiralles | Lachnospiraceae | Lachnobacterium |  |
| ***Lachnospiraceae*** | **1** | Bacillota | Clostridia | Lachnospiralles | Lachnospiraceae |  |  |
| ***Anaerococcus*** | **2** | Bacillota | Clostridia | Tissierellales | Peptoniphilaceae | Anaerococcus |  |
| ***Finegoldia*** | **2** | Bacillota | Clostridia | Tissierellales | Peptoniphilaceae | Finegoldia |  |
| ***Gallicola*** | **1** | Bacillota | Clostridia | Tissierellales | Peptoniphilaceae | Gallicola |  |
| ***Murdochiella*** | **1** | Bacillota | Clostridia | Tissierellales | Peptoniphilaceae | Murdochiella |  |
| ***Peptoniphilus*** | **3** | Bacillota | Clostridia | Tissierellales | Peptoniphilaceae | Peptoniphilus |  |
| ***Peptoniphilus lacrimalis*** | **1** | Bacillota | Clostridia | Tissierellales | Peptoniphilaceae | Peptoniphilus | lacrimalis |
| ***Phascolarctobacterium*** | **1** | Bacillota_C | Negativicutes | Acidaminococcales | Acidaminococcaceae | Phascolarctobacterium |  |
| ***Mitsuokella*** | **1** | Bacillota_C | Negativicutes | Selenomonadales | Selenomonadaceae | Mitsuokella |  |
| ***Negativicoccus succinivorans*** | **1** | Bacillota_C | Negativicutes | Veillonellales | Negativicoccaceae | Negativicoccus | succinivorans |
| ***Veillonella*** | **1** | Bacillota_C | Negativicutes | Veillonellales | Veillonellaceae | Veillonella |  |
| **Bacteroidota** | **1** | Bacteroidota |  |  |  |  |  |
| **Bacteroidia** | **1** | Bacteroidota | Bacteroidia |  |  |  |  |
| ***Prevotella*** | **2** | Bacteroidota | Bacteroidia | Bacteroidales | Bacteroidaceae | Prevotella |  |
| ***Prevotellaceae**** | **1** | Bacteroidota | Bacteroidia | Bacteroidales | Bacteroidaceae |  |  |
| ***Butyricimonas*** | **1** | Bacteroidota | Bacteroidia | Bacteroidales | Marinifilaceae | Butyricimonas |  |
| ***Odoribacter*** | **1** | Bacteroidota | Bacteroidia | Bacteroidales | Marinifilaceae | Odoribacter |  |
| ***Solitalea*** | **1** | Bacteroidota | Bacteroidia | Sphingobacteriales | Sphingobacteriaceae | Solitalea |  |
| ***Arcobacter*** | **1** | Campylobacterota | Campylobacteria | Campylobacterales | Arcobacteraceae | Arcobacter |  |
| ***Campylobacter*** | **1** | Campylobacterota | Campylobacteria | Campylobacterales | Campylobacteraceae | Campylobacter |  |
| **Campylobacterales** | **1** | Campylobacterota | Campylobacteria | Campylobacterales |  |  |  |
| ***Bilophila*** | **1** | Desulfobacterota | Desulfovibrionia | Desulfovibrionales | Desulfovibrionaceae | Bilophila |  |
| ***Desulfovibrio*** | **1** | Desulfobacterota | Desulfovibrionia | Desulfovibrionales | Desulfovibrionaceae | Desulfovibrio |  |
| ***Fusobacterium*** | **1** | Fusobacteriota | Fusobacteriia | Fusobacteriales | Fusobacteriaceae | Fusobacterium |  |
| **Nitrospirae*** | **1** | Nitrospirota |  |  |  |  |  |
| **Enhydrobacter*** | **1** | Pseudomonadota | Alphaproteobacteria | Reyranellales | Reyranellaceae | Reyranella |  |
| ***Agrobacterium*** | **1** | Pseudomonadota | Alphaproteobacteria | Rhizobiales | Rhizobiaceae | Agrobacterium |  |
| ***Rhodoplanes*** | **1** | Pseudomonadota | Alphaproteobacteria | Rhizobiales | Xanthobacteraceae | Rhodoplanes |  |
| ***Ramlibacter*** | **1** | Pseudomonadota | Gammaproteobacteria | Burkholderiales | Burkholderiaceae | Ramlibacter |  |
| ***Aeromonas*** | **1** | Pseudomonadota | Gammaproteobacteria | Enterobacterales | Aeromonadaceae | Aeromonas |  |
| ***Citrobacter*** | **1** | Pseudomonadota | Gammaproteobacteria | Enterobacterales | Enterobacteriaceae | Citrobacter |  |
| ***Enterobacter*** | **1** | Pseudomonadota | Gammaproteobacteria | Enterobacterales | Enterobacteriaceae | Enterobacter |  |
| ***Erwinia*** | **1** | Pseudomonadota | Gammaproteobacteria | Enterobacterales | Enterobacteriaceae | Erwinia |  |
| ***Klebsiella*** | **1** | Pseudomonadota | Gammaproteobacteria | Enterobacterales | Enterobacteriaceae | Klebsiella |  |
| ***Halomonas*** | **1** | Pseudomonadota | Gammaproteobacteria | Pseudomonadales | Halomonadaceae | Halomonas |  |
| ***Acinetobacter*** | **1** | Pseudomonadota | Gammaproteobacteria | Pseudomonadales | Moraxellaceae | Acinetobacter |  |
| ***Moraxellaceae*** | **1** | Pseudomonadota | Gammaproteobacteria | Pseudomonadales | Moraxellaceae |  |  |
| ***Pseudomonas*** | **1** | Pseudomonadota | Gammaproteobacteria | Pseudomonadales | Pseudomonaceae | Pseudomonas |  |
| **Pseudomonadales** | **1** | Pseudomonadota | Gammaproteobacteria | Pseudomonadales |  |  |  |
| ***Lysobacter*** | **1** | Pseudomonadota | Gammaproteobacteria | Xanthomonadales | Xanthomonadaceae | Lysobacter |  |
| ***Stenotrophomonas*** | **1** | Pseudomonadota | Gammaproteobacteria | Xanthomonadales | Xanthomonadaceae | Stenotrophomonas |  |
| **Synergistales** | **1** | Synergistota | Synergistia | Synergistales |  |  |  |
| **Synergistetes*** | **1** | Synergistota | Synergistia |  |  |  |  |
| ***Akkermansia muciniphila*** | **1** | Verrucomicrobiota | Verrucomicrobiae | Verrucomicrobiales | Akkermansiaceae | Akkermansia | muciphinila |
| *Higher abundance or more prevalent in urine of diabetics* |  |  |  |  |  |  |  |
| **Actinobacteriota** | **2** | Actinobacteriota |  |  |  |  |  |
| ***Mobiluncus curtisii*** | **1** | Actinomycetota | Actinomycetes | Actinomycetales | Actinomycetaceae | Mobiluncus | curtisii |
| ***Gardnerella vaginalis**** | **1** | Actinomycetota | Actinomycetes | Actinomycetales | Bifidobacteriaceae | Bifidobacterium | vaginale |
| ***Brevibacterium ravenspurgense*** | **1** | Actinomycetota | Actinomycetes | Actinomycetales | Brevibacteriaceae | Brevibacterium | ravenspurgense |
| ***Cellulosimicrobium*** | **1** | Actinomycetota | Actinomycetes | Actinomycetales | Cellulomonadaceae | Cellulosimicrobium |  |
| ***Microbacterium*** | **1** | Actinomycetota | Actinomycetes | Actinomycetales | Microbiobacteriaceae | Microbacterium |  |
| ***Micrococcus*** | **1** | Actinomycetota | Actinomycetes | Actinomycetales | Micrococcaceae | Micrococcus |  |
| ***Modestobacter*** | **1** | Actinomycetota | Actinomycetes | Mycobacteriales | Geodermatophilaceae | Modestobacter |  |
| ***Corynebacterium*** | **1** | Actinomycetota | Actinomycetes | Mycobacteriales | Mycobacteriaceae | Corynebacterium |  |
| ***Corynebacterium aurimucosum*** | **1** | Actinomycetota | Actinomycetes | Mycobacteriales | Mycobacteriaceae | Corynebacterium | aurimucosum |
| ***Corynebacterium coyleae*** | **1** | Actinomycetota | Actinomycetes | Mycobacteriales | Mycobacteriaceae | Corynebacterium | coyleae |
| ***Corynebacterium glucuronolyticum*** | **1** | Actinomycetota | Actinomycetes | Mycobacteriales | Mycobacteriaceae | Corynebacterium | glucuronolyticum |
| ***Propionibacterium*** | **1** | Actinomycetota | Actinomycetes | Propionibacteriales | Propionibacteriaceae | Propionibacterium |  |
| ***Collinsella*** | **1** | Actinomycetota | Coriobacteriia | Coriobacteriales | Coriobacteriaceae | Collinsella |  |
| ***Eggerthella*** | **1** | Actinomycetota | Coriobacteriia | Coriobacteriales | Eggerthellaceae | Eggerthella |  |
| ***Aerococcus*** | **1** | Bacillota | Bacilli | Lactobacillales | Aerococcaceae | Aerococcus |  |
| ***Aerococcus christensenii*** | **1** | Bacillota | Bacilli | Lactobacillales | Aerococcaceae | Aerococcus | christensenii |
| ***Enterococcus*** | **2** | Bacillota | Bacilli | Lactobacillales | Enterococcaceae | Enterococcus |  |
| ***Enterococcus faecalis*** | **2** | Bacillota | Bacilli | Lactobacillales | Enterococcaceae | Enterococcus | faecalis |
| ***Enterococcus faecium*** | **1** | Bacillota | Bacilli | Lactobacillales | Enterococcaceae | Enterococcus | faecium |
| ***Lactobacillus*** | **2** | Bacillota | Bacilli | Lactobacillales | Lactobacillaceae | Lactobacillus |  |
| ***Lactobacillus iners*** | **2** | Bacillota | Bacilli | Lactobacillales | Lactobacillaceae | Lactobacillus | iners |
| ***Turicibacter*** | **1** | Bacillota | Bacilli | MOL361 | Turicibacteraceae | Turicibacter |  |
| ***Anaerococcus hydrogenalis*** | **1** | Bacillota | Clostridia | Tissierellales | Peptoniphilaceae | Anaerococcus | hydrogenalis |
| ***Parvimonas*** | **1** | Bacillota | Clostridia | Tissierellales | Peptoniphilaceae | Parvimonas |  |
| ***Peptoniphilus grossensis*** | **1** | Bacillota | Clostridia | Tissierellales | Peptoniphilaceae | Peptoniphilus | grossensis |
| ***Veillonella atypica*** | **1** | Bacillota_C | Negativicutes | Veillonellales | Veillonellaceae | Veillonella | atypica |
| ***Veillonella montpellierensis*** | **1** | Bacillota_C | Negativicutes | Veillonellales | Veillonellaceae | Veillonella | montpellierensis |
| ***Allobaculum*** | **1** | Bacillota_I | Bacilli_A | Erysipelotrichales | Erysipelotrichaceae | Allobaculum |  |
| ***Prevotella bucalis*** | **1** | Bacteroidota | Bacteroidia | Bacteroidales | Bacteroidaceae | Prevotella | bucalis |
| ***Prevotella colorans*** | **1** | Bacteroidota | Bacteroidia | Bacteroidales | Bacteroidaceae | Prevotella | colorans |
| ***Porphyromonas*** | **1** | Bacteroidota | Bacteroidia | Bacteroidales | Porphyromonadaceae | Porphyromonas |  |
| ***Alistipes*** | **1** | Bacteroidota | Bacteroidia | Bacteroidales | Rikenellaceae | Alistipes |  |
| ***Flavobacteria**** | **1** | Bacteroidota | Bacteroidia |  |  |  |  |
| ***Flavobacteriales*** | **1** | Bacteroidota | Bacteroidia | Flavobacteriales |  |  |  |
| ***Chryseobacterium*** | **1** | Bacteroidota | Bacteroidia | Flavobacteriales | Weeksellaceae | Chryseobacterium |  |
| ***Bdellovibrio*** | **1** | Bdellovibrionota | Bdellovibrionia | Bdellovibrionales | Bdellovibrionaceae | Bdellovibrio |  |
| ***Campylobacter*** | **1** | Campylobacterota | Campylobacteria | Campylobacterales | Campylobacteraceae | Campylobacter | ureolyticus |
| ***Deinococcus*** | **1** | Deinococcota | Deinococci | Deinococcales | Deinococcaceae | Deinococcus |  |
| ***Anaeromyxobacter*** | **1** | Myxococcota | Myxococcia | Myxococcales | Anaeromyxobacteraceae | Anaeromyxobacter |  |
| **Pseudomonadota** | **1** | Pseudomonadota |  |  |  |  |  |
| ***Rubellimicrobium*** | **1** | Pseudomonadota | Alphaproteobacteria | Rhodobacterales | Rhodobacteraceae | Rubellimicrobium |  |
| ***Sphingomonas*** | **1** | Pseudomonadota | Alphaproteobacteria | Sphingomonadales | Sphingomonadaceae | Sphingomonas |  |
| ***Delftia**** | **1** | Pseudomonadota | Gammaproteobacteria | Burkholderiales | Burkholderiaceae | Comamonas |  |
| ***Hydrogenophaga*** | **1** | Pseudomonadota | Gammaproteobacteria | Burkholderiales | Burkholderiaceae | Hydrogenophaga |  |
| ***Methylophilus*** | **1** | Pseudomonadota | Gammaproteobacteria | Burkholderiales | Methylophilaceae | Methylophilus |  |
| ***Escherichia*** | **1** | Pseudomonadota | Gammaproteobacteria | Enterobacterales | Enterobacteriaceae | Escherichia |  |
| ***Escherichia coli*** | **1** | Pseudomonadota | Gammaproteobacteria | Enterobacterales | Enterobacteriaceae | Escherichia | coli |
| ***Escherichia-Shigella*** | **1** | Pseudomonadota | Gammaproteobacteria | Enterobacterales | Enterobacteriaceae | Escherichia-Shigella |  |
| ***Klebsiella*** | **1** | Pseudomonadota | Gammaproteobacteria | Enterobacterales | Enterobacteriaceae | Klebsiella |  |
| **Haemophilus** | **1** | Pseudomonadota | Gammaproteobacteria | Enterobacterales | Pasteurellaceae | Haemophilus |  |
| ***Stenotrophomonas*** | **1** | Pseudomonadota | Gammaproteobacteria | Xanthomonadales | Xanthomonadaceae | Stenotrophomonas |  |
